# BatchDTA: Implicit batch alignment enhances deep learning-based drug-target affinity estimation

**DOI:** 10.1101/2021.11.23.469641

**Authors:** Hongyu Luo, Yingfei Xiang, Xiaomin Fang, Wei Lin, Fan Wang, Hua Wu, Haifeng Wang

**Affiliations:** Baidu Inc., 518000, Shenzhen, China; Baidu Inc., 100000, Beijing, China

**Keywords:** Drug-target affinity, Deep neural network, Batch alignment, Batch effects

## Abstract

Candidate compounds with high binding affinities toward a target protein are likely to be developed as drugs. Deep neural networks (DNNs) have attracted increasing attention for drug-target affinity (DTA) estimation owning to their efficiency. However, the negative impact of batch effects caused by measure metrics, system technologies, and other assay information is seldom discussed when training a DNN model for DTA. Suffering from the data deviation caused by batch effects, the DNN models can only be trained on a small amount of “clean” data. Thus, it is challenging for them to provide precise and consistent estimations. We design a batch-sensitive training framework, namely BatchDTA, to train the DNN models. BatchDTA implicitly aligns multiple batches toward the same protein, alleviating the impact of the batch effects on the DNN models. Extensive experiments demonstrate that BatchDTA facilitates four mainstream DNN models to enhance the ability and robustness on multiple DTA datasets. The average concordance index (CI) of the DNN models achieves a relative improvement of 4.0%. BatchDTA can also be applied to the fused data collected from multiple sources to achieve further improvement.

## Introduction

Evaluation of drug-target affinity (DTA) (also known as receptor-ligand affinity) is one of the fundamental tasks for drug discovery. The drug-target binding affinity indicates the strength of binding interaction between the drug (ligand/compound) and target (receptor/protein). Laboratory experiments [27], e.g., in vivo and in vitro experiments, are usually involved in measuring the affinities between the protein to be studied and the candidate compounds. The compounds with the highest measured affinities are screened for further validation and could be future drugs. Since laboratory experiments are laborious, expensive, and time-consuming, some advanced studies employ efficient machine learning methods to estimate binding affinities, accelerating drug discovery. Machine learning methods [5, 26, 7, 6, 14], especially deep neural networks (DNNs) [31, 35, 32, 28, 34, 22, 30, 18, 29, 16] have gained increasing attention of many scholars, which utilize the affinity data collected from laboratory experiments to infer the affinities of other protein-compound interactions.

The Batch effects, as common effects in data from high-throughput laboratory experiments, have been widely discussed in many previous studies [24, 8]. This paper defines the batch effects as the systematic batch variation caused by differences in measure metrics, system technologies, laboratory conditions, reagents, and other assay information. Many published studies [1, 3] have shown that the systematic variation in experimentally collected data may lead to incorrect biological conclusions. Lots of techniques [23, 12, 38] have attempted to detect and adjust batch effects in laboratory experiments, but the negative impact of batch effects is still barely discussed in the field of DTA estimation, especially DTA estimation through DNN models.

The model parameters of the mainstream DNN models for DTA [31, 32, 22, 28, 18, 16] are usually optimized through the pointwise training framework, which minimizes the distance between the estimated affinity and the ground-truth affinity for each protein-compound interaction. Due to the lack of consideration of batch effects in model training, the DNN models trained by the pointwise framework is difficult to fully exploit the existing affinity data from multiple sources, measured by various techniques and systems, and could lead to the following issues:

First, the assays of affinity evaluation may adopt various metrics, e.g., dissociation constant (*K*_*D*_), inhibition constant (*K*_*I*_), half-maximal inhibitory concentration (*IC*_50_), and half-maximal effective concentration (*EC*_50_), to measure the binding affinities. The affinity values evaluated through different metrics are usually incomparable (refer to Appendix B). A DNN model trained by the pointwise framework is confused by the multifarious incomparable “affinities” from multiple assays/batches of the same protein-compound interaction, thus struggling to provide a consistent estimation. Previous academic works tend to train a DNN model on the data points measured by the same metric to get around the metric incomparability issue. Nevertheless, the data points with respect to a specific metric are insufficient to train a capable DNN model, usually with numerous parameters and thus requiring lots of training data points.

Second, even though only utilizing those data points of a specific metric to train a model, the trained model still suffers from variance caused by other assay information. For example, some metrics, e.g., *IC*_50_ and *EC*_50_, are highly assay-specific [19], and the measured affinities under different assay settings are incomparable. The DNN model is confounded by the multifarious “affinities” from multiple assays of the same protein-compound interaction, thus struggling to provide consistent estimates.

Consequently, the batch effects caused by various metrics, assay settings, and other factors obstruct advanced DNN models from fully exploiting the collected data points or achieving more satisfactory precision. It is attractive to take advantage of fuse multi-source data points with various assay settings and align these data points to further enhance the DNN model’s ability.

To facilitate the effectiveness of the DNN models for DTA, we attempted to alleviate the negative impact of batch effects through a batch-sensitive training framework, namely BatchDTA. BatchDTA learns the ranking orders of the comparable protein-compound interactions in the same batch. In this way, the interactions of a protein from multiple batches can be implicitly aligned by taking those interactions that simultaneously appeared in multiple batches as the reference for comparison. Extensive experiments are conducted to verify that through training with BatchDTA framework, multiple mainstream DNN models, including DeepDTA [31], GraphDTA [28], and MolTrans [16], can achieve higher concordance indexes (CIs) and lower deviation. We also visualize a case to demonstrate that BatchDTA can successfully learned the ranking orders of the compounds from different batches through the reference interaction. Furthermore, BatchDTA is applied to utilize fused data points measured by multiple metrics from multiple sources as the training data. The concordance indexes (CIs) [11] of the DNN models are promoted in most cases, exhibiting the potential practical value of BatchDTA.

Our main contributions can be summarized as follows:

1. We utilize a batch-sensitive training framework, called BatchDTA, to alleviate the harmful influence of the batch effects for DTA prediction.
2. BatchDTA aims to implicitly align multiple batches for the same target protein by learning the ranking orders of the interactions within a batch.
3. Extensive experiments demonstrate that BatchDTA can facilitate the precision and robustness of multiple mainstream DNN models in multiple situations.

## Materials and Methods

### Problem Formulation

Advanced proposed DNN models for DTA utilize the pointwise training framework, which regards DTA estimation as a classification or regression problem. Since the goal of the DTA estimation task is locating those compounds with higher affinities toward a target protein, we regard the task of DTA estimation as a ranking problem instead. More concretely, a ranking task of DTA is defined as a batch (or an assay) with batch *b* = (*p, 𝒞*), containing a protein *p* and a set of the candidate compounds *𝒞*. The set of candidate compounds is defined as 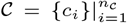, where *n*_*𝒞*_ represents the number of candidate compounds in the corresponding batch. We expect to screen out the most promising compounds in *𝒞* as the potential drugs for target protein *p* for batch *b*. Usually, hundred to tens of thousands of batches can be collected from a data source, 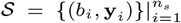, where *n*_*s*_ represents the number of batches in data source *𝒮*. The set of the corresponding labels, i.e., collected affinities, of the candidate compounds *C* in batch *b* is denoted by 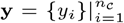. Additionally, we attempt to simultaneously exploit the data from various data sources 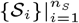 to train the DNN models for DTA to test the prospect of data fusion, where *n*_*S*_ denotes the number of data sources.

### Implicit Batch Alignment

We attempt to align a target protein’s corresponding protein-compound interactions from multiple batches. Those interactions are implicitly aligned through the reference interactions that simultaneously appear in multiple batches. The demonstration of implicit batch alignment is shown in Fig. 1, and the details will be introduced in the following.

**Fig. 1.**
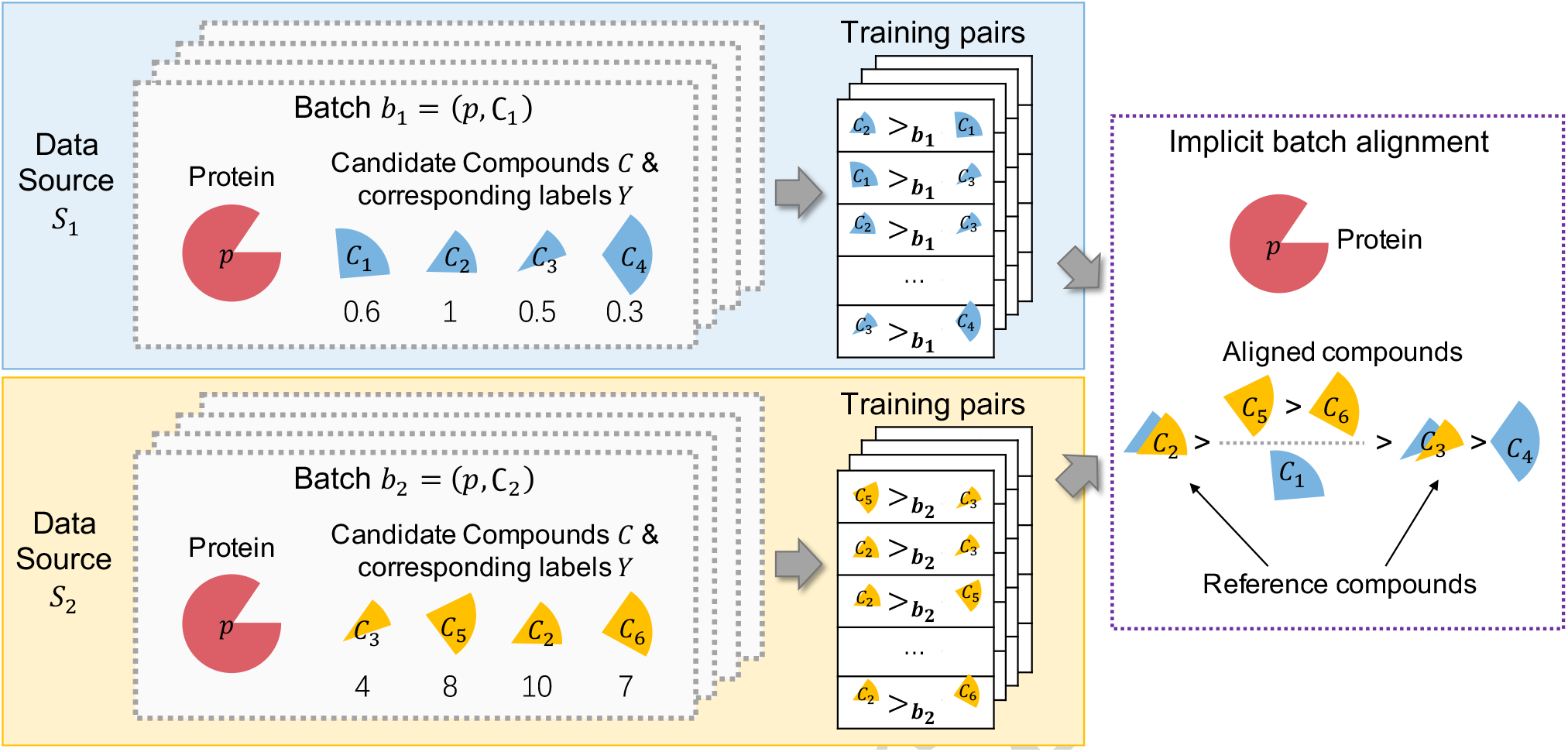
Implicit batch alignment. Compound pairs for training are generated from each batch. The compounds in multiple batches for the same protein are implicitly aligned through the compounds in both batches. (Notched circles represent proteins and sectors of various colors represent compounds)

We first define the ranking order of two compounds concerning a batch. Since two interactions from different batches may target different proteins or their corresponding affinities may be evaluated under different assay settings, the ranking orders of most compound pairs are valueless. Thus, we concentrate on learning the ranking order of two candidate compounds *c*_*i*_ and *c*_*j*_ for a given batch *b* = (*p, 𝒞*). To formalize, we introduce a new operator *>*_*b*_ for comparison of two compounds. Formula *c*_*i*_ *>*_*b*_ *c*_*j*_ indicates the affinity of compound *c*_*i*_ measured by a assay is significantly larger than that of compound *c*_*j*_ for the same batch *b*. Since affinities measured by the laboratory assays are usually noisy, a hyper-parameter, i.e., deviation *E*, is introduced to determine whether the two measured affinities differ significantly. More concretely, formula *c*_*i*_ *>*_*b*_ *c*_*j*_ indicates that *y*_*i*_ *> y*_*j*_ + *E*, where *y*_*i*_ and *y*_*j*_ denotes the assay-measured affinities of compound *c*_*i*_ and compound *c*_*j*_ in the compound set *C* of the batch *b* = (*p, 𝒞*). Then, for a batch *b* = (*p, 𝒞*), we can theoretically generate 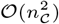 training pairs of compounds: 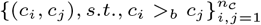. We define the training dataset as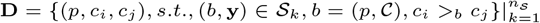. Pair (*c*_*i*_, *c*_*j*_) is called a training pair as demonstrated on the left of Fig. 1, and triple (*p, c*_*i*_, *c*_*j*_) is called a training sample.

Then, we introduce reference compounds to align the interactions from different batches. A reference compound is defined as a compound that appears in multiple batches toward the same protein. Given a protein *p*, there might be multiple batches studying that protein. We assume both batch *b*_1_ = (*p, 𝒞*_1_) and batch *b*_2_ = (*p, 𝒞*_2_) study protein *p*. A reference compound *c*^*ref*^ should satisfy that *c*^*ref*^ *∈𝒞*_1_ *∧ c*^*ref*^ *∈𝒞*_2_. For batch *b*_1_ = (*p, 𝒞*_1_), we denote the compounds in *𝒞*_1_ with the corresponding ground-truth affinities smaller than that of the reference compound *c*^*ref*^ as 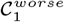. For batch *b*_2_ = (*p, 𝒞* _2_), the compounds in *𝒞* _2_ with the corresponding ground-truth affinities larger than that of the reference compound *c*^*ref*^ as 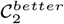. It is likely the affinity of compound 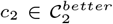 is larger than that of compound 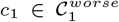 In this way, the compounds from different batches can be implicitly aligned. A toy case is exhibited in Fig. 1. Both the batch *b*_1_ = (*p, 𝒞* _1_) in data source *S*_1_ and the batch *b*_2_ = (*p, 𝒞* _2_) in data source *S*_2_ study the protein *p*, where *𝒞* _1_ = *{c*_1_, *c*_2_, *c*_3_, *c*_4_*}* and *C*_2_ = *{c*_3_, *c*_5_, *c*_2_, *c*_6_*}*. Due to the batch effects, the affinity values collected from various sources are incomparable (e.g., from 0.3 to 1.0 in batch *b*_1_ and from 4 to 10 in batch *b*_2_). Training pairs (compound pairs) are extracted from each batch based on the ranking order of the compounds. The training pairs from different batches with respect to protein *p* are collected. As compound *c*_3_ appear in *𝒞* _1_ and *𝒞* _2_, compound *c*_3_ is taken as a reference compound. Though the reference compound *c*_3_, compound *c*_4_ from batch *b*_1_, compounds *c*_5_, and *c*_6_ from batch *b*_2_ become comparable.

Since compounds from different batches are comparable through implicit batch alignment, we can effortlessly fuse the data points from multiple sources to train a more capable model for DTA.

### Learning the Ranking Orders

Many studies utilized deep neural networks (DNNs) to predict the affinity between proteins and compounds, including DeepDTA [31], GraphDTA [28], and MolTrans [16]. We train these DNN models by learning the ranking orders of the training pairs.

A typical DNN model for DTA prediction consists of three components: Protein Encoder, Compound Encoder, and Interaction Estimator, as shown on the left of Fig. 2.

**Fig. 2.**
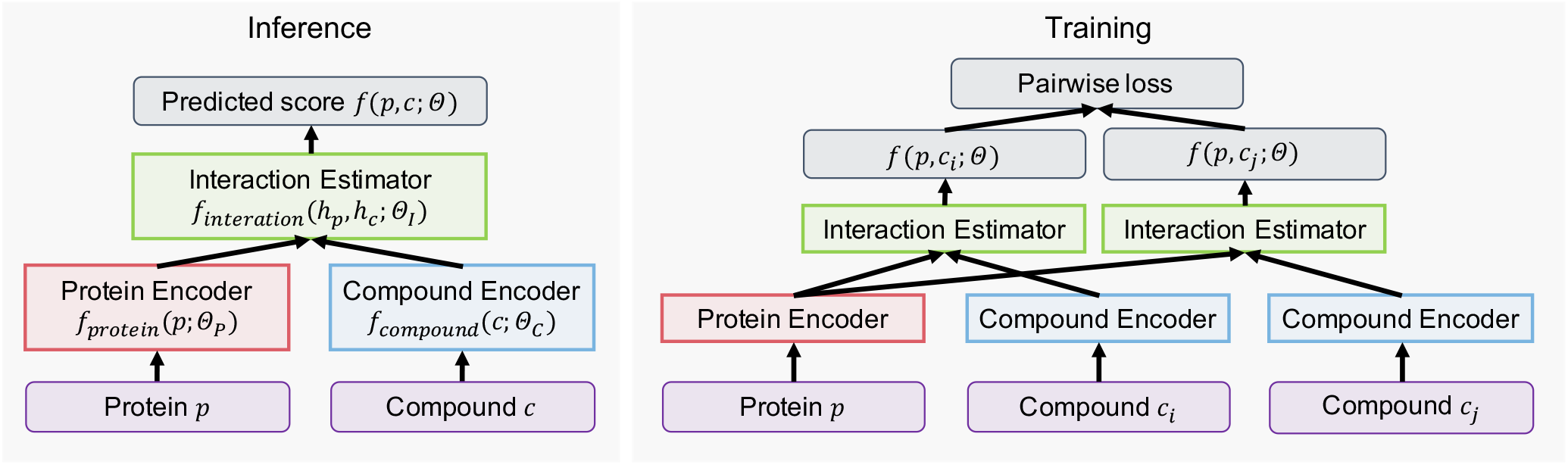
Inference of a typical DNN model *f* (*p, c*; Θ) for DTA and the training framework that trains the DNN model *f* (*p, c*; Θ) through learning the ranking orders of the candidate compounds.

- **Protein Encoder**. The primary structure of a protein is described by an amino acid sequence. A Protein Encoder applies various kinds of sequence-based neural networks, e.g., Convolutional Neural Networks (CNNs) [21, 20] and Transformers [37], to produce the representation vectors of the proteins. We formalize the Protein Encoder as a function *f*_*protein*_(*p*; Θ_*P*_), taking a protein *p* as input, where Θ_*p*_ represents the corresponding model parameters. The representation vector of a protein *p* is defined as *h*_*p*_ = *f*_*protein*_(*p*; Θ_*P*_).
- **Compound Encoder**. A Compound Encoder typically generates the compounds’ representation vectors through molecular fingerprints (e.g., ECFP [33] and MACCS [10]), sequence-based representation methods (e.g., LSTMs [15] and Transformers), or graph-based representation methods (e.g., GNNs). We describe the Compound Encoder as a function *f*_*compound*_(*c*; Θ_*C*_), taking a compound *c* as input, where Θ_*C*_ represents the model parameters needed to be optimized. Then, the representation vector of a compound *c* can be written as *h*_*c*_ = *f*_*compound*_(*c*; Θ_*C*_).
- **Interaction Estimator**. The Interaction Estimator estimates the interaction score between a protein *p* and a compound *c* in accordance to a function *f*_*interation*_(*h*_*p*_, *h*_*c*_; Θ_*I*_), where Θ_*I*_ denotes the function’s learnable parameters. Function *f*_*interation*_(*h*_*p*_, *h*_*c*_; Θ_*I*_) takes the representation vectors *h*_*c*_ and *h*_*p*_ as inputs and outputs an estimated affinity score.

The parameters of whole model *f* (*p, c*; Θ) can be formulated as Eq. 1, combining *f*_*protein*_(*p*; Θ_*P*_), *f*_*compound*_(*c*; Θ_*C*_), and

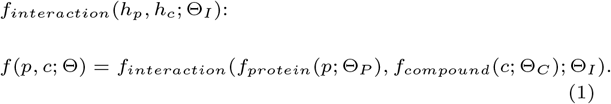

where Θ = {Θ_*C*_, Θ_*P*_, Θ_*I*_ } are the model parameters need to be optimized. The candidate compounds are ranked in accordance with the affinities estimated by the DNN model *f* (*p, c*; Θ), and the compounds with the highest estimated affinities are selected as promising drugs for further validation.

Advanced studies optimize the model parameters Θ to minimize the absolute error between the assay-measured affinity and the model-estimated affinity of a protein-compound pair. In contrast to the previously applied training framework (pointwise training), we adopt a pairwise training framework to optimize the ranking orders of the candidate compounds of a given batch, as shown on the right of Fig. 2. More concretely, the pairwise training framework takes the sample (*p, c*_*i*_, *c*_*j*_) *∈* **D** as input to learn the order of compounds *c*_*i*_ and *c*_*j*_ with respect to protein *p*. Following the classical pairwise ranking method RankNet [4], we regard the pairwise order learning problem as a binary classification task, where the cross-entropy is used as the loss function for the sample (*p, c*_*i*_, *c*_*j*_) *∈* **D**:

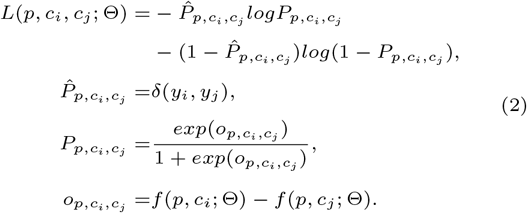

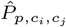 indicates whether the ground-truth affinity score of compound *c*_*i*_, i.e., *y*_*p,c*_*i* is larger than that of compound *c*_*j*_, i.e., *y*_*p,c*_*j*. *δ*(*x, y*) is an indicator function that *δ*(*x, y*) = 1 if *x > y*, and *δ*(*x, y*) = 0 if *x ≤ y*. Besides, *P*_*p,c*_*i*_,*c*_*j* represents the model-estimated probability whether the affinity of compound *c*_*i*_ is larger than that of compound *c*_*j*_. *P*_*p,c*_*i*_,*c*_*j* applies Sigmoid function [13] on *o*_*p,c*_*i*_,*c*_*j*, i.e., the difference between the estimated affinities *f* (*p, c*_*i*_; Θ) and *f* (*p, c*_*j*_ ; Θ). The cross-entropy between the ground-truth probability 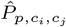 and the estimated probability *P*_*p,c*_*i*_,*c*_*j* is minimized to learn the model parameters of the DNN model *f* (*p, c*; Θ). As an affinity pair (*p, c*_*i*_, *c*_*j*_) *∈* **D** satisfies *c*_*i*_ *>*_*b*_ *c*_*j*_ and *y*_*i*_ *> y*_*j*_, 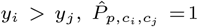, the loss function for a sample (*p, c*_*i*_, *c*_*j*_) in Eq. 2 can be simplified as

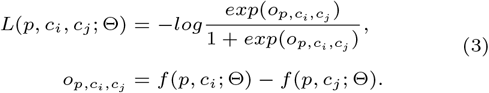

Then, the overall loss function of dataset **D** is defined as

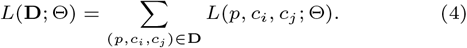

We minimize the loss function *L*(**D**; Θ) to optimize the model parameters of *f* (*p, c*; Θ).

As we mentioned above, we can theoretically produce 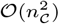 samples for a batch *b* = (*p, 𝒞*). In order to balance the effects of different batches for training, we randomly sample *n*_*sample*_ samples for each compound within a batch *b* at each iteration, where *N* is the sampling times. In this way, for each batch, only *O*(*n*_*sample*_ · *n*_*𝒞*_) samples are exploited for training at each iteration.

## Results

We use the pointwise framework and BatchDTA to train multiple advanced DNN models, respectively, to observe the effect of BatchDTA.

### Datasets

The DNN models are trained on three DTA datasets, including BindingDB [25], Davis [9], and KIBA [36]. The BindingDB dataset collected affinities with various metrics (*K*_*D*_, *K*_*I*_, *IC*_50_, and *EC*_50_) and other assay information from multiple data sources. The affinities in the Davis dataset are measured by *K*_*D*_, while those in the KIBA dataset are indicated by the KIBA score that integrates multiple bioactivity types, i.e., *K*_*D*_, *K*_*I*_, and *IC*_50_. These datasets are processed to filter out the invalid protein-compound interactions (refer to Appendix. A) for data processing).

Previous DTA studies [28, 18, 32] randomly split the protein-compound interactions of a dataset into a training set and a test set. However, random splitting over-simplifies the task of DTA. Due to multiple batches, a target protein in the test set could have already been observed in the training set, resulting in information leakage and model overfitting. Therefore, we split the datasets on the basis of batches and ensure that the proteins observed in the training set will not appear in the test set, which is more in line with the real-world applications. For each dataset, we strive to identify the batches based on the available assay information recorded in the corresponding dataset. Since the batch IDs are not recorded in the datasets, we can not completely guarantee the accuracy of batch identification (refer to Appendix. A for batch identification of each dataset). The statistics of the datasets are shown in Tab. 2.

### DNN Models for DTA

We compare two training frameworks, *Pointwise* and *BatchDTA*, by applying them to three previously proposed DNN models for DTA:

- **DeepDTA** [31] employs three-layers Convolutional Neural Networks (CNNs) as Protein Encoder and Compound Encoder to encode the protein sequences and the compound SMILES strings, respectively. Then, for the Interaction Estimator, the encoded protein and compound are concatenated to predict the affinity score.
- **GraphDTA** [28] regards each compound as a graph and attempts several GNNs, such as GIN, GAT, GCN, and GAT-GCN, as the Compound Encoders to represent the compounds. In the meantime, GraphDTA regards each protein as a sequence and adopts CNNs as the Protein Encoder to encode the proteins. The Interaction Estimator is the same as DeepDTA’s. In this paper, we implement two versions of GraphDTA, denoted by GraphDTA(GCN) and GraphDTA(GATGCN), respectively.
- **MolTrans** [16] decomposes the compounds’ SMILES strings and the proteins’ acid amino sequences into high-frequency sub-sequences. Then, it applies Transformers as the Compound Encoder and Protein Encoder to obtain the augmented representation with the chemical semantics. MolTrans uses the outer-product operator and CNN blocks as Interaction Estimator to capture the high-order interaction between the compounds and proteins.

For all the DNN models, we use the hyper-parameters suggested by the corresponding papers.

### Training and Evaluation Settings

We follow the settings of the previous works to implement *Pointwise* framework, where the mean squared error (MSE) between the models’ predicted values and the ground-truth affinities are taken as the loss function to optimize the model parameters. For the *BatchDTA* framework, we apply grid search to search the hyper-parameters, including sample times, learning rate, and batch size. The details of hyper-parameter settings are described in Appendix C.1.

Concordance Index (CI) [11] is used to evaluate the performance of *Pointwise* and *BatchDTA* training frameworks in terms of batches. The CI for a given batch *b* = (*p, C*) is defined as:

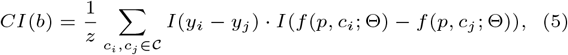

where 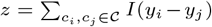, and *I*(.) is an indicator function. *I*(*x*) = 1 if *x > ϵ*^*\**^, and *I*(*x*) = 0 otherwise. *ϵ*^*\**^ is used to identify the compound pairs with large gap of their corresponding affinities. *ϵ*^*\**^ set to be 0 by default unless otherwise specified. Then, we define the overall CI for all the test batches 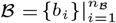 by summarizing *CI*(*b*), where *n*_*B*_ is the number of test batches. The overall CI is formalized as

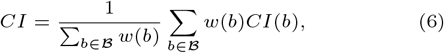

where *w*(*b*) = *n*_*𝒞*_ for task *b* = (*p, 𝒞*) is introduced to balance the impact of different batches.

### Overall Performance

To verify that implicit batch alignment can alleviate the negative impact of batch effects, we trained multiple previously proposed DNN models for DTA through the *Pointwise* and *BatchDTA* training frameworks, respectively. We expect a DNN model trained by *BatchDTA* framework can achieve further improvement compared with that trained by *Pointwise* framework.

Database BindingDB records the affinities of millions of protein-compound interactions measured by various metrics. Most data points in BindingDB are measured by metrics inhibition constant (*K*_*I*_) and half-maximal inhibitory concentration (*IC*_50_). Two hundred thousand protein-compound interactions from ten thousand batches are measured by *K*_*I*_, while six hundred thousand interactions from twenty thousand batches are measured by *IC*_50_ (refer to Table 2 for statistics of the datasets). Since the assay information, such as temperature, pH degree, and competitive substrate, affect the values of *K*_*I*_ and *IC*_50_, the data points measured by *K*_*I*_ and *IC*_50_ are suitable for verifying whether *BatchDTA* can effectively mitigate the impact of batch effects. The dataset of each metric is divided into the training set, validation set, and test set by 8:1:1. A model is trained on the training set with the best epoch selected by the validation set. For each experimental setting, we trained each model five times and evaluated the CIs on all the compound pairs (*ϵ*^*\**^ = 0) as well as the confident compound pairs (*ϵ*^*\**^ = 0.6) in the test set. Confident compound pairs are introduced to diminish the impact of the experimental variance for the evaluation. We eliminated the compound pairs with small affinity gaps from all the compound pairs, and the remaining pairs are called confident compound pairs (refer to Appendix C.2) for details of confident compound pairs generation).

Fig. 3 shows the CIs of four DNN models trained by *Pointwise* framework and *BatchDTA* framework. We can draw the following conclusions:

**Fig. 3.**
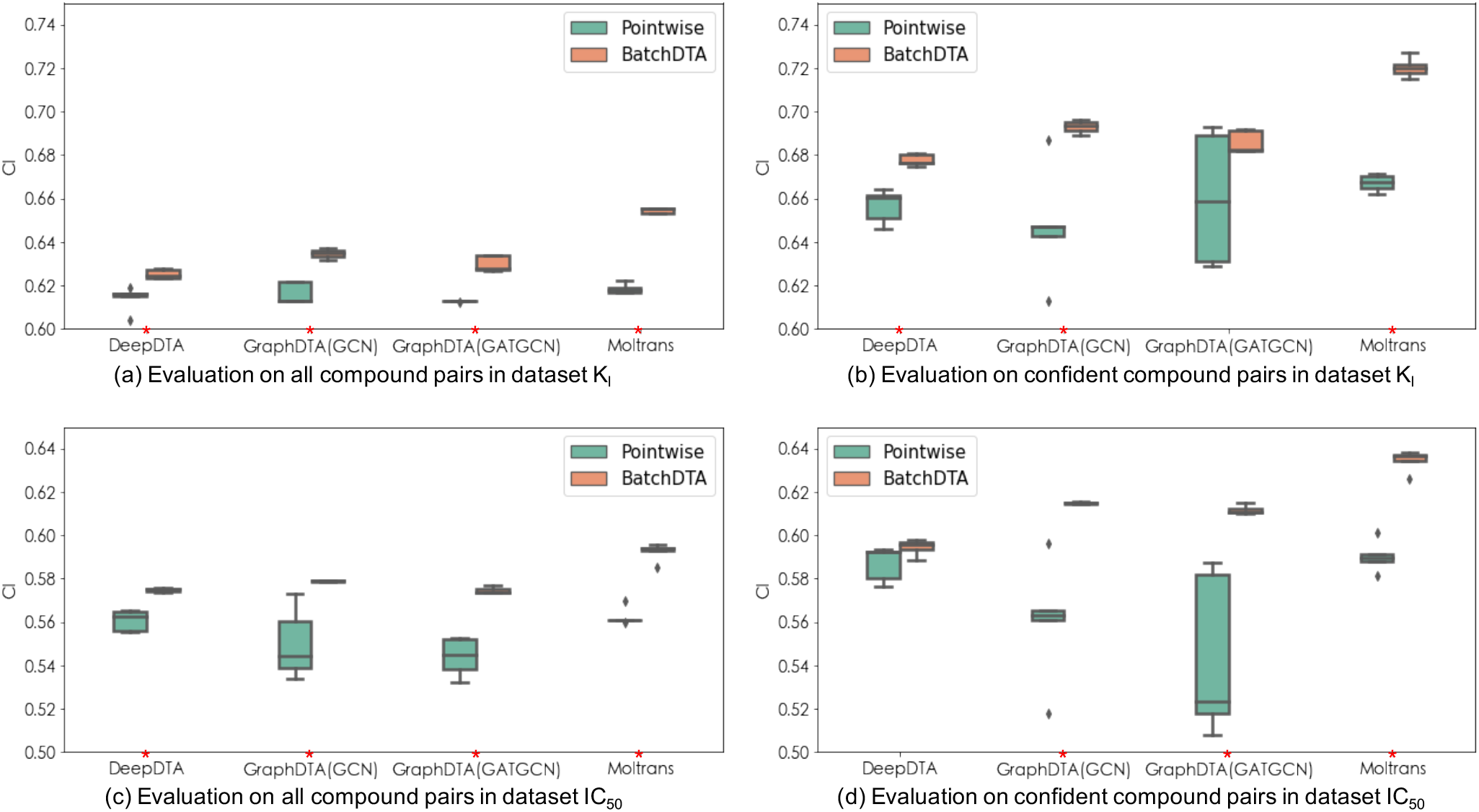
Comparing the CIs of the DNN models trained by *Pointwise* framework and *BatchDTA* framework on database BindingDB under different settings. (A red star stands for statistical significance with P-value less than 0.05 for a two-tailed test.)

- Since *BatchDTA* is a batch-sensitive training framework, it can make more rational use of the collected data points than the *Pointwise* framework, which could further enhance the ability and robustness of a DNN model for DTA. *BatchDTA* framework significantly outperforms *Pointwise* framework under 14/16 settings. Besides, in most cases, the standard deviation of CIs of the model trained by the *Pointwise* framework is greater than that by the *BatchDTA* framework. We suspect that the conflicting data between batches cause the larger variance of the model trained by the *Pointwise* framework.
- For each DNN model, the relative improvement of *BatchDTA* compared with *Pointwise* is higher when evaluating on the confident compound pairs than on all the compound pairs. Since the gaps between the measured affinities of some compounds in a batch are not significant, excluding those incomparable compound pairs in the evaluation can highlight the advantages of *BatchDTA*. For example, for *K*_*I*_, *BatchDTA* achieves an average relative improvement of 3.2% on all the compound pairs, while 5.5% on the confident compound pairs. For *IC*_50_, the average relative improvement is 4.7% for all the compound pairs and 7.8% for confident compound pairs.
- Compared with *K*_*I*_, *IC*_50_ is also affected by the concentration of enzyme and substrate, and thus the systematic variance of the assays measured by *IC*_50_ is larger than that measured by *K*_*I*_. The larger variance makes a DNN model more difficult to provide accurate estimations on the *IC*_50_ dataset, but we can observe that all the DNN models trained by *BatchDTA* can still achieve further improvement.

### Case Study

We further explore whether *BatchDTA* can successfully align and learn the ranking orders of the compounds from multiple batches by case study. We selected a case of two batches toward the same protein from the training set of *IC*_50_ with a reference compound appearing in both batches. The reference compound is expected to implicitly align the compounds from two batches. In Fig. 4, we visualize three matrices with each element representing the affinity gap between a compound in Batch 1 and a compound in Batch 2. The affinity gaps are normalized, and the darker the color in the matrices, the greater the affinity gap. More concretely, we sorted compounds in Batch 1 and Batch 2 according to the ground-truth affinities, respectively. For each matrix, each column stands for a compound from Batch 1, each row stands for a compound for Batch 2, and each element stands for a compound pair with its color indicating the affinity gap. The three matrices describe the gaps between the ground-truth affinities collected in the dataset, the gaps between the predicted affinities of the MolTrans model trained by the *Pointwise* framework, the gaps between the predicted affinities of the MolTrans model trained by *BatchDTA* framework, respectively.

**Fig. 4.**
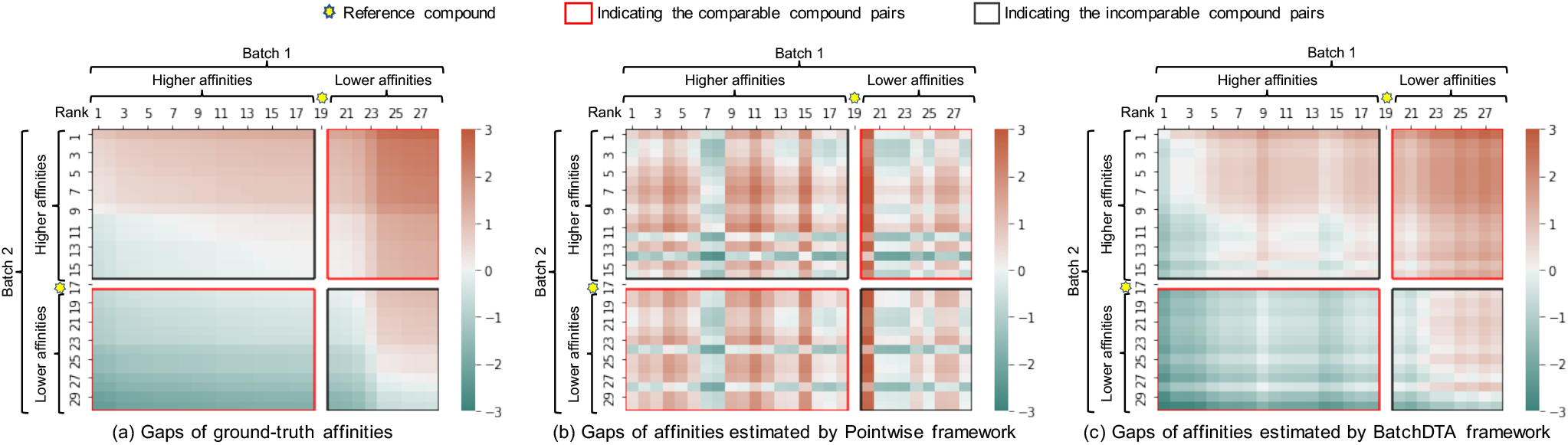
Case study analyzing whether *Pointwise* framework and *BatchDTA* framework can learn the ranking orders of the compound from two batches for the same protein.

In general, the color pattern of the gaps of affinities estimated by *BatchDTA* framework (Fig. 4(c)) is similar to that of the gaps of ground-truth affinities (Fig. 4(a)). The colors in both the ground-truth matrix and *BatchDTA* matrix transition smoothly, with dark red at the upper right, dark green at the lower left, and white at the diagonal. The consistent color patterns reveal that *BatchDTA* can well learn the ranking order of a compound pair with one compound in Batch 1 and the other in Batch 2. In contrast, the color pattern of *Pointwise* is chaotic, which is distinct from that of the ground-truth. More specifically, the grids (compound pairs) in a matrix can be divided into two portions: the comparable compound pairs indicated by the red boxes (lower left and upper right) and the incomparable compound pairs indicated by the black boxes (lower right and upper left). For example, the compounds with higher affinities than the reference compound in Batch 1 (the first 18 columns) should rank ahead of those with lower affinities than the reference compound in Batch 2 (the last 13 rows). Thus, the corresponding compound pairs indicated by the red box at the lower left are regarded as comparable compound pairs. The model trained by *BatchDTA* can infer the compounds’ ranking order with a high certainty (indicated by darker color) for most of the comparable compound pairs. For the incomparable compound pairs, although the model trained by *BatchDTA* can not determine some compounds’ ranking order, e.g., the ranking order of the first compound in Batch 1 and the first compound in Batch 2, the overall ranking predictions are consistent with those of the ground-truth.

### Fusing Data

The *BatchDTA* framework can also be applied to the fused data points collected from multiple sources. It is attractive to take advantage of fused data points to train the DNN models for DTA. Therefore, we explore the practicality of *BatchDTA* in two data fusion scenarios: fusing data measured with multiple metrics and fusing data from multiple open-source datasets.

### Fusing Data Points Measured by Multiple Metrics

We utilized the database BindingDB and trained the DNN models on the fusing data measured with multiple metrics. Each DNN model is trained on four datasets, i.e., *K*_*I*_, *K*_*I*_ &*K*_*D*_, *K*_*I*_ &*K*_*D&*_*IC*_50_, and *K*_*I*_ &*K*_*D&*_*IC*_50_&*EC*_50_, respectively by *BatchDTA* framework, where operator & denotes fusing the data points measured by different metrics. The trained DNN models are evaluated on the test set of *K*_*I*_. We consider the correlations between the metrics (refer to Appendix B) when designing the experiments. Since the metrics with higher correlations to *K*_*I*_ are more likely to provide advantages to the DNN models that only trained on the data points of *K*_*I*_, we first integrated the data points of *K*_*D*_ into the training set, then those of *IC*_50_, and finally those of *EC*_50_. Each experiment is run five times, and the performance of the DNN models trained on the fused data points measured by multiple metrics is shown in Fig. 5. We can observe the following phenomena: (1) By adding the data points of *K*_*D*_ to the training set, all the DNN models gain significant performance improvement (average relative improvement of 1.21%). Since *K*_*I*_ and *K*_*D*_ have a strong correlation (Pearson r=0.83), the data points of *K*_*D*_ potentially augment valuable information. (2) Due to the higher systematic deviations of the data measured by *IC*_50_ and *EC*_50_, the correlation between *K*_*I*_ and *IC*_50_ as well as the correlation between *K*_*I*_ and *EC*_50_ are relatively low (Pearson r=0.74 and 0.64, respectively). Thus, fusing the data points of *IC*_50_ and *EC*_50_ can not further significantly raise the accuracy of the DNN models. These phenomena enlighten us that integrating high correlation data into the training set could improve the effectiveness of existing DNN models on DTA.

**Fig. 5.**
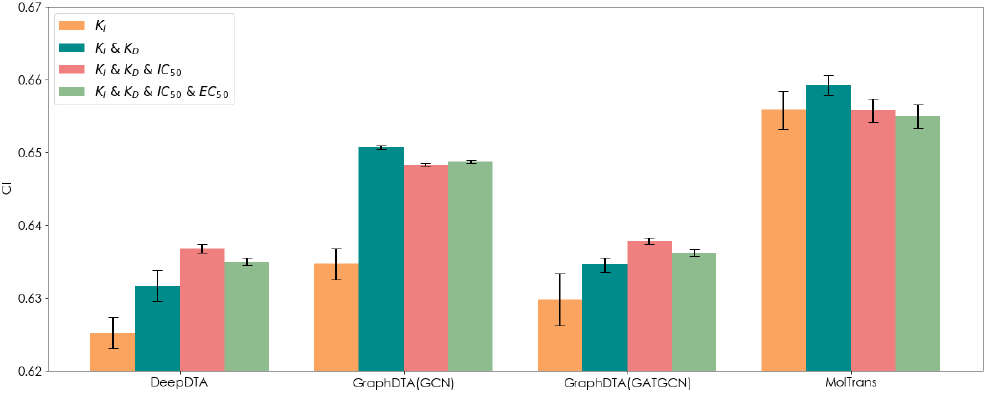
Exploring the performance of the DNN models trained on the fused data points measured by multiple metrics, i.e., *K*_*I*_, *K*_*D*_, *IC*_50_, and *EC*_50_ by *BatchDTA* framework.

### Fusing Data Points from Multiple Datasets

Datasets Davis and KIBA are utilized to explore the potential of *BatchDTA* when training the DNN models on the fused data points collected from multiple datasets. Since these two datasets contain only hundreds of batches (refer to Table 2 in Appendix), we perform 5-fold cross-validation [2] to reduce the evaluation deviation. When evaluating the DNN models on the Davis dataset, we compare three training methods: *Pointwise (Davis)* and *BatchDTA (Davis)* that only utilize the Davis dataset for training, as well as *BatchDTA (Davis + KIBA)* that utilize both the datasets Davis and KIBA for training. The setting is similar when evaluating the DNN models on the KIBA dataset. Fig. 6 exhibits the comparison of the DNN models trained on the fused data points collected from multiple datasets. As we expected, the CIs of the DNN models trained on only one dataset by *BatchDTA* framework surpass those by *Pointwise* framework, which is consistent with the evaluation results of Fig. 3 on database BindingDB. Furthermore, by combining the datasets Davis and KIBA for training, the DNN models trained by *BatchDTA* achieve better performance in general. The better results of the fused datasets illustrate that as a DNN model usually requires lots of training data, the additional data points for training are likely to benefit the ability of the DNN models for DTA.

**Fig. 6.**
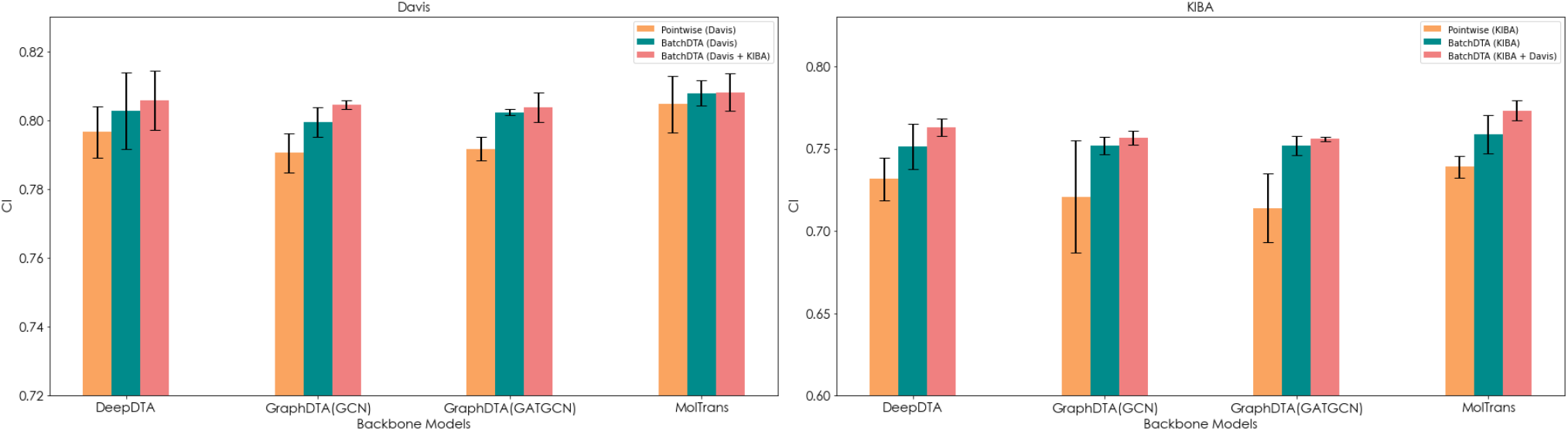
Exploring the performance of the DNN models trained on the fused data points collected from multiple datasets, i.e., Davis and KIBA, by *BatchDTA* framework.

## Discussion

Many published literature have discussed the impact of batch effects, but little work has been done to explore the batch effects on the DNN models for DTA. Table 1 reports the data analysis of the database BindingDB. We attempt to analyze the impact scope of batch effects for DTA through the dimensions of proteins and batches (the first two rows in Table 1) and investigate which group of the impacted data points (the last row in Table 1) can be effectively taken advantage of by BatchDTA to achieve better results. From the table, we can observe: (1) About 40% of proteins simultaneously appear in multiple batches, and the collected affinities of corresponding data points could be inconsistent. A batch-sensitive training framework is demanded to alleviate the negative impact of those inconsistent data points. (2) There are 43.3% to 55.7% of batches with interactions appearing in another batch for each metric. That means about half of the batches can be implicitly aligned through those interactions (reference compounds), and BatchDTA has the opportunity to model those batches better. (3) We extracted the compound pairs that appeared in multiple batches against the same protein from the datasets and found that the ranking orders of a small number of compound pairs (7.1% to 14.6%) were inconsistent. We suspect the inconsistency is caused by the assay errors, batch identification errors, biological variance, and other factors. Employing BatchDTA to train DNN models can not overcome the impact of those noisy data points. Fortunately, the ranking orders of only less than 1% of all the compound pairs are inconsistent.

**Table 1.** Data analysis of dataset BindingDB.

| | $K_D$ | $K_I$ | $IC_{50}$ | $EC_{50}$ | All |
| --- | --- | --- | --- | --- | --- |
| Proteins appearing in multiple batches | 40.0% | 42.2% | 40.5% | 36.1% | 44.0% |
| Batches with interactions appearing in another batch | 43.3% | 55.7% | 48.5% | 48.6% | 57.9% |
| Inconsistent ranking orders | 12.4% | 8.1% | 7.1% | 14.6% | 13.9% |

We further investigate the impact of the quality of batch identification. Fig. 7 compares the effects of two batch identification methods. For the first method *Batch*, we try our best to distinguish different batches according to the fields recorded in the database. For the second method *Protein*, all the data points with respect to the same protein are grouped in the same batch. Thus, the quality of method *Batch* is higher than that of method *Protein* for batch identification. The results of dataset *K*_*I*_ (Fig. 7(a)) illustrate that the quality of batch identification plays a critical role in the effectiveness of BatchDTA. Besides, for dataset *IC*_50_, the CIs of methods *Batch* and *Protein* are comparable. Since *IC*_50_ is affected by more factors as mentioned, but those factors are not recorded in the database, we can not precisely identify the batches from the database. We believe that the DNN models trained by BatchDTA will benefit from more detailed assay information.

**Fig. 7.**
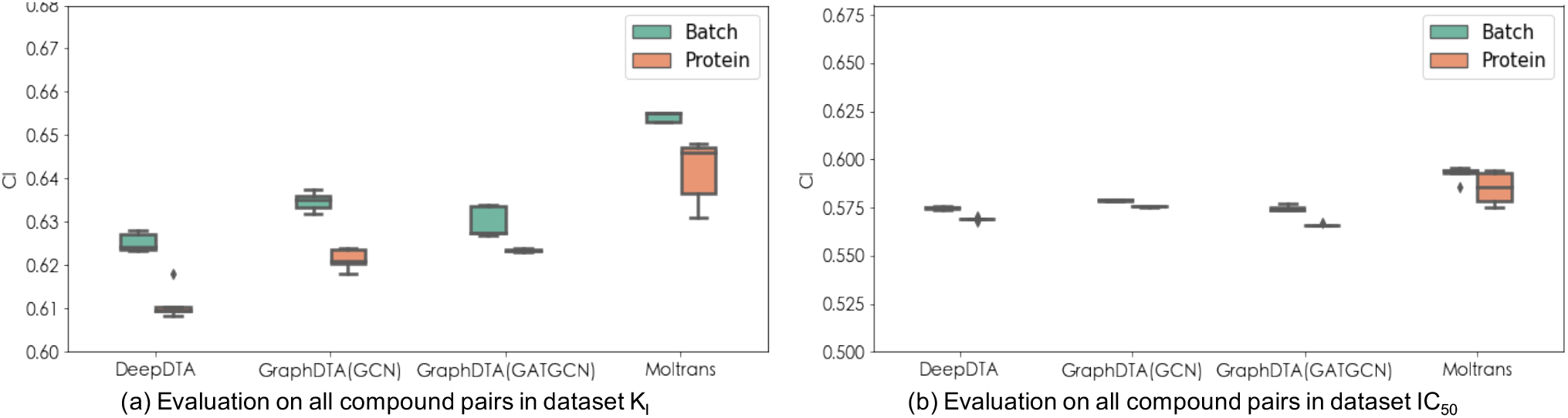
Impact of the quality of batch identification methods.

## Conclusions

Many advanced studies have applied deep learning to drug-target affinity (DTA) estimation to screen the promising compounds toward the given target proteins. Existing studies focus more on designing DNN model architectures for DTA. We argue that a batch-sensitive training framework, i.e., BatchDTA, can further enhance the accuracy and robustness of the previously proposed DNN models, weakening the influence of batch effects (systematic variance). Each batch is regarded as a ranking task. BatchDTA implicitly aligns the corresponding compounds toward the same protein from multiple batches by learning the compounds’ ranking orders in each batch. Extensive experimental results demonstrate that four advanced DNN models trained by BatchDTA can achieve significant improvement on three datasets for DTA. BatchDTA can also be applied to train the DNN models on fused data collected from multiple sources to further enhance the models’ precision. We believe our work could inspire researchers working on applying machine learning methods to drug discovery.

## Appendix: Datasets and Data Process

We describe the details of the data process in this section, and the data statistics of the processed datasets are shown in Table 2.

**Table 2.** Statistics of the datasets.

| Dataset | Metric | #Compound | #Protein | #Interaction | #Batch |
| --- | --- | --- | --- | --- | --- |
| BindingDB | $K_D$ | 6,704 | 587 | 31,239 | 1,070 |
| | $K_I$ | 126,122 | 1,225 | 249,598 | 11,292 |
| | $IC_{50}$ | 407,462 | 2,582 | 653,344 | 20,041 |
| | $EC_{50}$ | 75,142 | 699 | 105,364 | 3,143 |
| Davis | $K_D$ | 68 | 442 | 30,056 | 442 |
| KIBA | KIBA score | 2,111 | 229 | 118,254 | 229 |

### BindingDB

The BindingDB database with the version of May 2021 contains 2,221,487 compound-protein interactions, including 7,965 proteins and 963,425 compounds.

First, we cleaned up the raw data from BindingDB by the following steps: (1) Keep compound-protein interactions measured by metrics *K*_*D*_, *K*_*I*_, *IC*_50_ and *EC*_50_). (2) Remove the recorded affinities with ‘*>*‘ or ‘*<*‘. (3) Rescale the affinities larger than 10,000 to be 10,000. (4) Drop out the duplicate records. (5) Remove the compounds with illegal SMILES strings that can not be processed by the Cheminformatics software RDKit (https://rdkit.org). (6) Remove the proteins that can not be converted to FASTA format (https://zhanggroup.org/FASTA/).

Second, we identify the batches from the cleaned data: (1) Multiple fields in BindingDB, including ‘BindingDB Target Chain Sequence’, ‘pH’, ‘Temp (C)’, ‘Curation/DataSource’, ‘Article DOI’, ‘PMID’, ‘PubChem AID’, ‘Patent Number’, ‘Authors’, and ‘Institution’, are considered to identify batches. (2) Remove the batches with less than ten candidate compounds.

### Davis and KIBA

We followed the previous work [31] to process Davis and KIBA datasets, and the implementation of which is at https://github.com/hkmztrk/DeepDTA/tree/master/data. Then, the batches are identified according to ‘Gene ID’ for the Davis dataset and ‘Uniprot ID’ for the KIBA dataset.

## Appendix Relations between Metrics of Binding Affinities

Dissociation constant (*K*_*D*_), inhibition constant (*K*_*I*_), half-maximal inhibitory concentration (*IC*_50_), and half-maximal effective concentration (*EC*_50_) are mainly used for measuring binding affinities. Both *K*_*D*_ and *K*_*I*_ are used to indicate the dissociation equilibrium constant, i.e., the reverse of the association constant, for the dissociated components. *K*_*D*_ is a more general term, while *K*_*I*_ is more specifically used under enzyme-inhibitor complex. *IC*_50_ is the inhibitory concentration, measured by an inhibitor with the response (or binding) reduced by half. *EC*_50_ stands for effective concentration of an inhibitor with half-maximal response. Comparing with *K*_*I*_ and *K*_*D*_, *IC*_50_ and *EC*_50_ are influenced by more assay information, such as substrate concentration [17].

In particular, through Cheng-Prusoff equation, *K*_*I*_ can be converted from *IC*_50_ [19]:

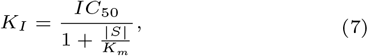

where |*S*| is the substrate concentration, and *K*_*m*_ is the Michaelis-Menten constant of the substrate. Thus, *IC*_50_ is assay-specific, and the *IC*_50_ values under different assay information are incomparable.

In Section. 3.6.1, we evaluate the DNN models that are trained on fused data measured by multiple metrics on the test set of *K*_*I*_. The data points are added gradually according to the Pearson correlation of the corresponding metrics to *K*_*I*_. The Pearson correlations are shown in Fig. 8. We first added the data points of *K*_*D*_, then *IC*_50_, and finally *EC*_50_.

**Fig. 8.**
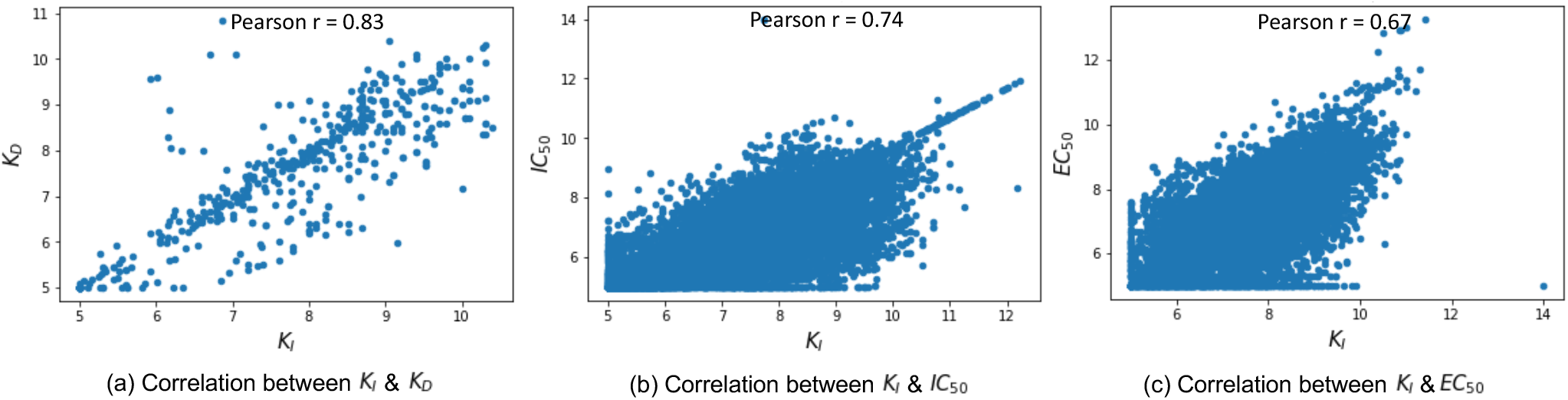
Pearson correlations between *K*_*I*_ and other metrics.

## Appendix Experimental Settings

### Hyper-parameters of BatchDTA

Following the previous works [31, 14, 28, 18], we normalize the original affinities as labels for training. Take *K*_*D*_ for example. We use *pK*_*D*_ values as labels to train the DNN models:

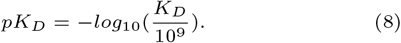

*K*_*I*_, *IC*_50_ and *EC*_50_ and KIBA score are normalized in the same way.

We used Adam optimizer and applied grid search to explore the best hyper-parameters for each DNN model on each dataset.

For BindingDB database, the candidate settings of each hyper-parameter are shown in Table 3.

**Table 3.** Candidate settings of hyper-parameters for BindingDB.

| Hyper-parameter | Candidate settings |
| --- | --- |
| deviation $\epsilon$ | {0.2, 0.5, 1} |
| $n_{sample}$ for $K_I$ | 10 |
| $n_{sample}$ for $K_D$ | {1, 3, 5} |
| $n_{sample}$ for $IC_{50}$ | {0.2, 0.5, 1} |
| $n_{sample}$ for $EC_{50}$ | {0.1, 0.2, 0.5} |
| learning rate | $\{1 \times 10^{-3}, 5 \times 10^{-4}\}$ |
| batch size | {256, 512} |

For Davis and KIBA datasets, the candidate settings of the hyper-parameters for searching are shown in Table 4.

**Table 4.** Candidate settings of hyper-parameters for Davis and KIBA.

| Hyper-parameter | Candidate settings |
| --- | --- |
| deviation $\epsilon$ | {0.2, 0.5, 1} |
| $n_{sample}$ | {0.5, 1, 3, 5, 10} |
| learning rate | $\{1 \times 10^{-3}, 5 \times 10^{-4}\}$ |
| batch size | {256, 512} |

All the models are trained by NVIDIA Tesla V100 GPUs. Since BatchDTA is a training framework, it would not change the number of model parameters and the time complexity for training.

### Generation of Confident Compound Pairs

To generate the confident compound pairs for BindingDB database, we eliminated the compound pairs with small affinity gaps from all the compound pairs. The probability of the compound pairs with affinity gaps smaller than *ϵ*^*\**^ is shown in Fig. 9. When *ϵ*^*\**^ = 0.6, about 50% compound pairs are eliminated.

**Fig. 9.**
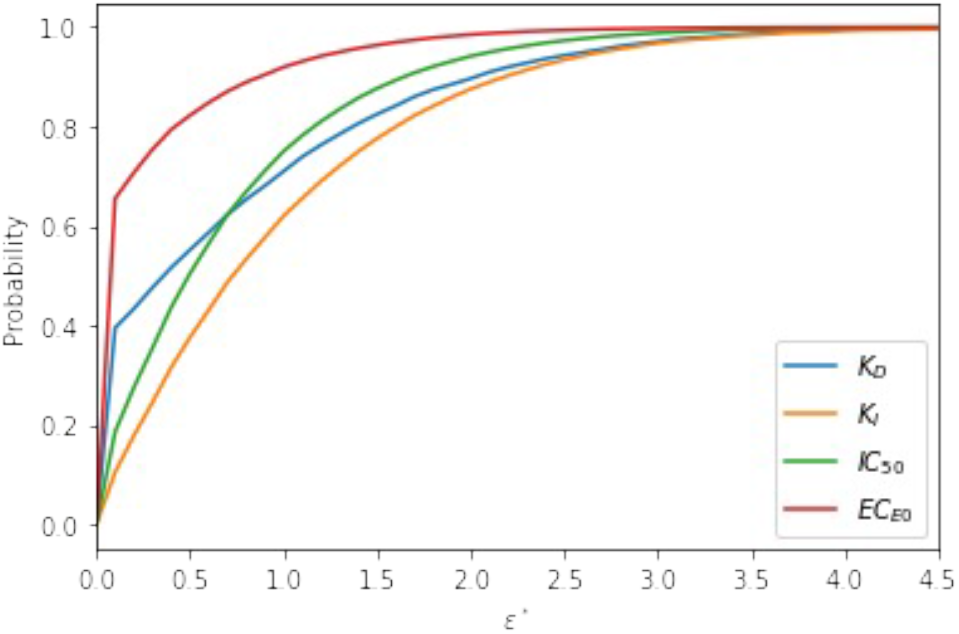
Probability of the compound pairs with affinity gaps smaller than *ϵ*.

## Appendix Detailed Results

### Overall Performance

Table 5 shows the detailed results corresponding to Fig.3. The DNN models trained by *BatchDTA* framework significantly outperform those trained by *Pointwise* framework in most cases.

**Table 5.**
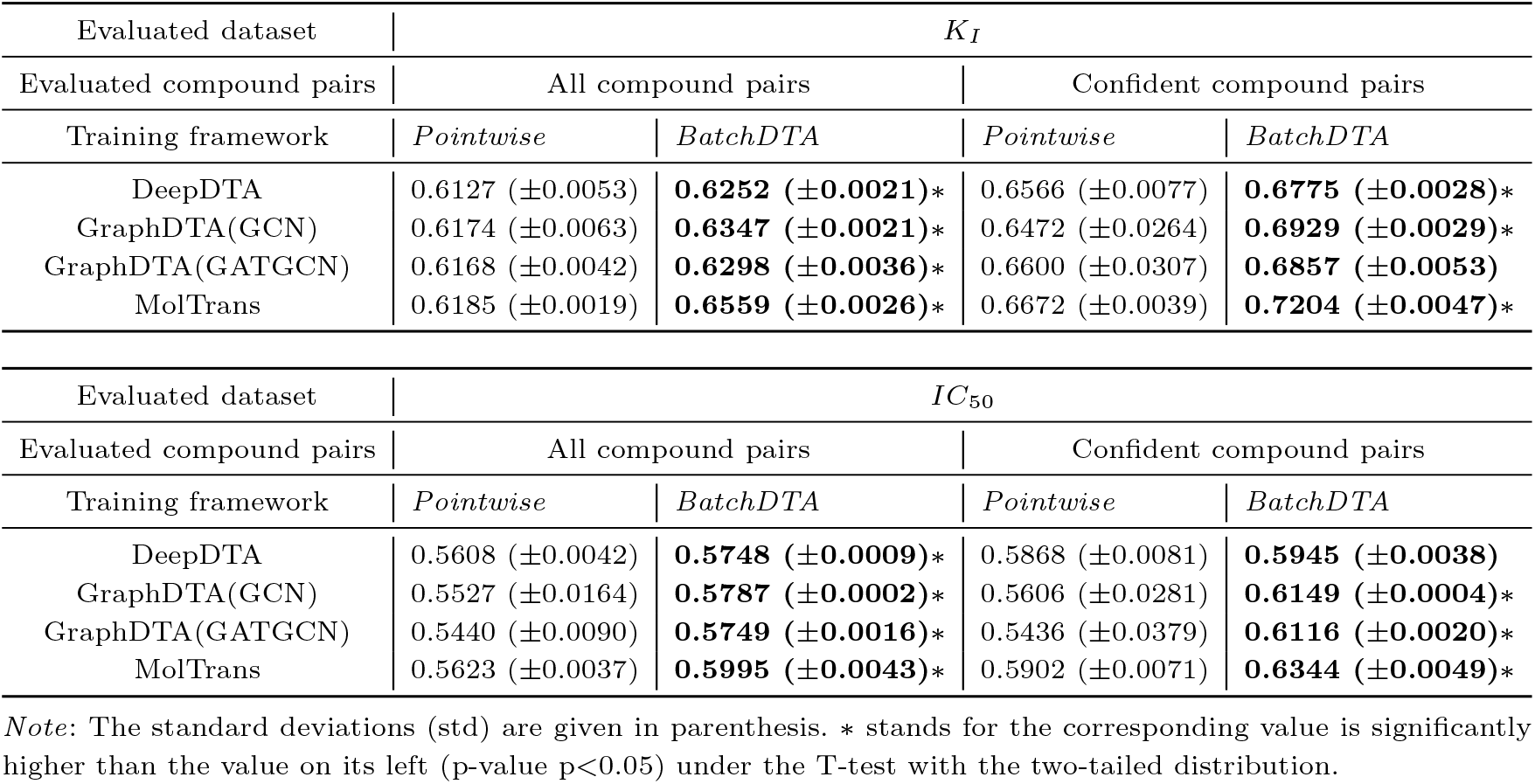
The detailed results corresponding to Fig. 3

### Fusing Data Points Measured by Multiple Metrics

Table 6 shows the detailed results corresponding to Fig.5. All the models achieve significant improvement when fusing the data points of *K*_*D*_. Two models, DeepDTA and GraphDTA(GATGCN), gain significant improvement when fusing the data points of *IC*_50_, but no model can get further improvement when fusing the data points of *EC*_50_. These results indicate that the data points with higher correlations play a more important role for data argumentation.

**Table 6.** The detailed results corresponding to Fig. 5h

| Training framework | BatchDTA |  |  |  |
| --- | --- | --- | --- | --- |
| Training dataset | $K_I$ | $K_I \& K_D$ | $K_I \& K_D \& IC_{50}$ | $K_I \& K_D \& IC_{50} \& EC_{50}$ |
| DeepDTA | 0.6252 ($\pm 0.0021$) | 0.6317 ($\pm 0.0021$)* | **0.6368 ($\pm 0.0005$)\*** | 0.6350 ($\pm 0.0003$) |
| GraphDTA(GCN) | 0.6347 ($\pm 0.0021$) | **0.6507 ($\pm 0.0003$)\*** | 0.6483 ($\pm 0.0019$) | 0.6488 ($\pm 0.0002$) |
| GraphDTA(GATGCN) | 0.6298 ($\pm 0.0036$) | 0.6346 ($\pm 0.0010$)* | **0.6377 ($\pm 0.0065$)\*** | 0.6363 ($\pm 0.0006$) |
| MolTrans | 0.6559 ($\pm 0.0026$) | **0.6595 ($\pm 0.0004$)\*** | 0.6558 ($\pm 0.0016$) | 0.6550 ($\pm 0.0015$) |

*Note:* The standard deviations (std) are given in parenthesis. \* stands for the corresponding value is significantly higher than the value on its left (p-value $p < 0.05$ ) under the T-test with the two-tailed distribution.

### Fusing Data Points from Multiple Datasets

Table 7 shows the detailed results corresponding to Fig.6. As we expected, the CIs of the DNN models trained by *BatchDTA* framework significantly surpass those trained by *Pointwise* framework in most cases, consistent with the results in the BindingDB database (refer to Fig. 3). Besides, training on the fusing data points from Davis and KIBA allows the DNN models to achieve better performance than training on a single dataset. Especially when evaluating the models on the KIBA dataset, three models gain significant improvement. Since the KIBA dataset contains only two hundred batches, we suspect that combining four hundred batches from the Davis dataset enhances the richness of batches.

**Table 7.**
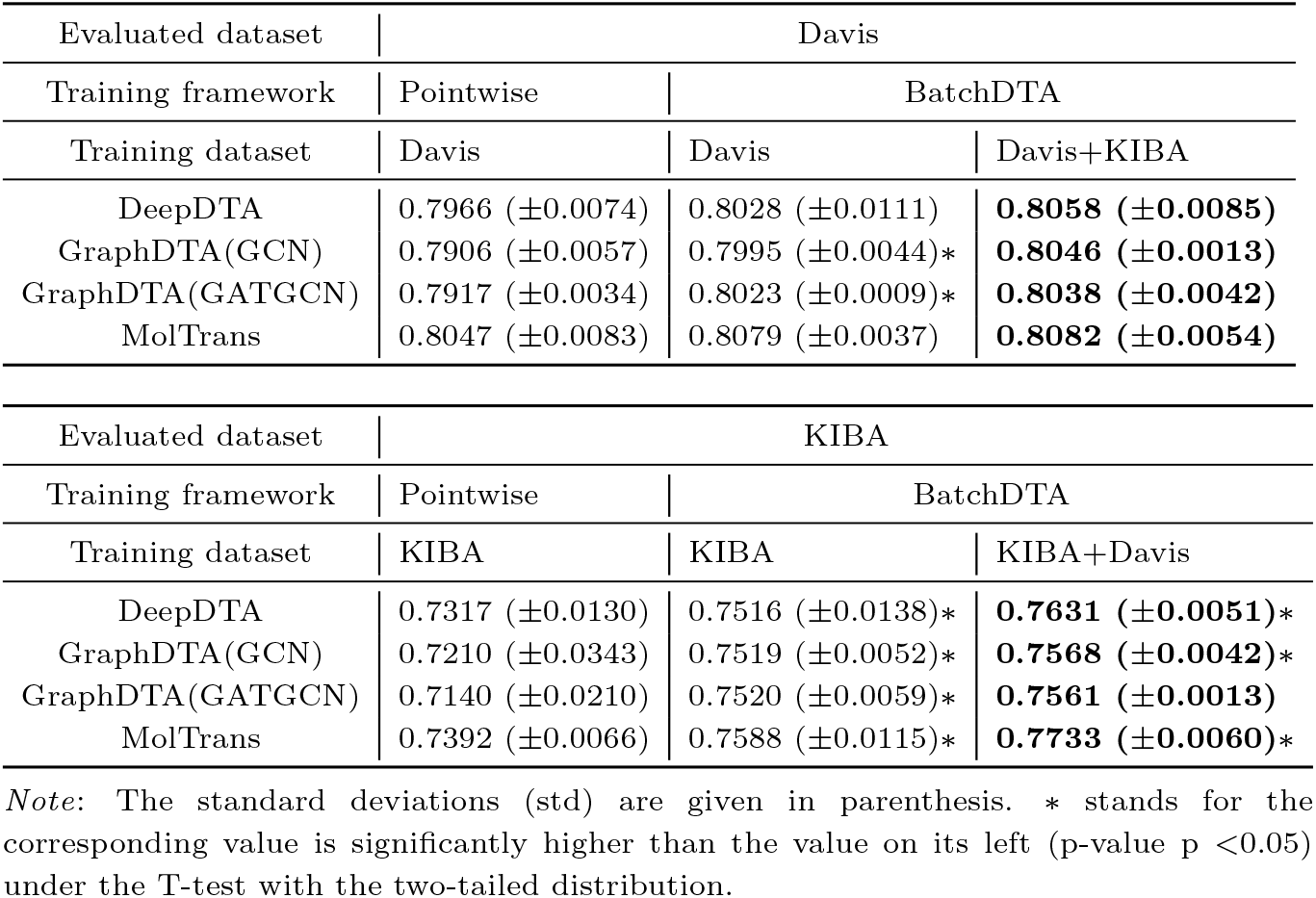
The detailed results corresponding to Fig. 6

## Code Availability

The code is freely available at GitHub https://github.com/PaddlePaddle/PaddleHelix/tree/dev/apps/drug_target_interaction/batchdta to allow replication of the results.

## Data Availability

The raw datasets, BindingDB, Davis, and KIBA are collected from https://www.bindingdb.org/bind/index.jsp, http://staff.cs.utu.fi/~aatapa/data/DrugTarget/ and https://jcheminf.biomedcentral.com/articles/10.1186/s13321-017-0209-z. The processed data is available at GitHub https://github.com/PaddlePaddle/PaddleHelix/tree/dev/apps/drug_target_interaction/batchdta to allow replication of the results.

## Author Contributions Statement

XM.F., F.W., H.W., and HF.W. led the research. HY.L, YF.X, XM.F., and F.W. conceived the experiments. HY.L. YF.X, and W.L.conducted the experiments. LY.L, XM.F., and YF.X. wrote the manuscript.

## Acknowledgments

The authors thank the anonymous reviewers for their valuable suggestions.

